# SCExecute: cell barcode-stratified analyses of scRNA-seq data

**DOI:** 10.1101/2022.03.27.485988

**Authors:** Nathan Edwards, Christian Dillard, NM Prashant, Hongyu Liu, Mia Yang, Evgenia Ulianova, Anelia Horvath

## Abstract

**Motivation:** In single-cell RNA-sequencing (scRNA-seq) data, stratification of sequencing reads by cellular barcode is necessary to study cell-specific features. However, apart from gene expression, the analyses of cell-specific features are not supported by available tools that are designed for bulk RNA-Seq data.

**Results:** We introduce a tool – SCExecute – which executes a user-provided command on barcode-stratified, extracted on-the-fly, single cell binary alignment map (scBAM) files. SCExecute extracts the cell barcode from aligned, pooled single-cell sequencing data. The user-specified command option executes all the commands defined in the session from monolithic programs and multi-command shell-scripts to complex shell-based pipelines. The execution can be further restricted to barcodes or/and genomic regions of interest. We demonstrate SCExecute with two popular variant callers - GATK and Strelka2 – combined with modules for bam file manipulation and variant filtering, to detect single cell-specific expressed Single Nucleotide Variants (sceSNVs) from droplet scRNA-seq data (10X Genomics Chromium System).

**Conclusion:** SCExecute facilitates custom cell-level analyses on barcoded scRNA-seq data using currently available tools and provides an effective solution for studying low (cellular) frequency transcriptome features.

**Availability:** SCExecute is implemented in Python3 using the PySAM package and distributed for Linux and Python environments from https://github.com/HorvathLab/NGS/tree/master/SCExecute.

## Introduction

In single-cell RNA-sequencing (scRNA-seq) data cell barcodes are used to extract cell-specific sequencing reads and assess cell-level features. Methods for cell-specific scRNA-seq analysis have been focused on gene expression, where popular approaches – such as STARsolo and CellRanger – integrate alignment with concurrent barcode demultiplexing and the assignment of read counts to genes (Kaminow *et al*., 2021; Tran *et al*., 2019). Additional cell-level transcriptome feature analyses – for example, expressed genetic variation, allele-specific expression and splicing – are now beginning to emerge, creating new knowledge, and demonstrating the substantial information content of scRNA-seq data (Liu *et al*., 2021; Prashant *et al*., 2020; N. M. Prashant *et al*., 2021; La Manno *et al*., 2018). These types of analyses can benefit from broader cell-level read-stratifying methods.

To aid custom cell-level analyses of scRNA-seq data, we have developed SCExecute – a software that executes a desired command or a set of commands on each barcode-stratified single-cell binary alignment map (scBAM) file. SCExecute generates scBAMs on-the-fly by extracting barcodes from the QNAME field or tags of barcoded, aggregated single-cell alignments, produced by widely used scRNA-seq tools, such as CellRanger, STARsolo, and UMI-tools (Smith *et al*., 2017). ScBAMs and the respective downstream analyses, can be restricted to a user-specified list of barcodes, such as the filtered list of cell-barcodes generated by STARsolo, or user-selected barcodes of interest. The user-specified command option executes all the commands defined in the session from monolithic programs and multi-command shell-scripts to complex shell-based pipelines. ScBAMs can optionally be filtered for reads aligned to genomic regions of interest.

## Software description

### Implementation

SCExecute is modeled after the classic Linux tools find and xargs, which execute a command repeatedly on a collection of files. Instead of generating scBAMs for all cellular-barcodes first, and only then executing a command on the scBAMs, SCExecute interleaves the generation of a batch of scBAMs and the execution of the command – avoiding operating system limits, such as on the number of open file handles – and enabling maximum throughput on multiprocessor systems. Users can specify the number of commands to run at once and adjust the batch size to ensure that all processors are kept busy. User commands are executed in the user’s command-line shell – so monolithic programs and multi-command shell-scripts, and complex commands including pipes, list operators, and standard input, output, and error redirection work as usual.

SCExecute provides explicit configuration for alignments barcoded using CellRanger, STARsolo, and UMItools. Users can also configure novel cellular barcode extraction logic based on BAM file tags, or values in the QNAME field (or, to its analogous counterpart – sequence identifier of the fastq files) and delimited tokens or regular expressions. The generated scBAMs filename and other execution specific parameters are inserted into the user-command before execution, using the same syntax as find and xargs, to specify the location of the generated filename (“{}”). In addition, the cell-barcode is available for substitution using “{BARCODE}” and the name of the input BAM file containing pooled scRNA-Seq alignments, without its path or the “.bam” extension, is available as “{BAMBASE}”. If no replacement tokens are used in the user-command template, the generated scBAMs file is provided as the last argument to the command. Users can also specify similarly templated filenames for generated scBAMs filenames, working directories for executing the command, and filenames for capturing standard output and/or standard error. The generated scBAMs retain the header fields and sorted order of the pooled scRNA-Seq BAM file provided as input. For debugging or focused analyses, SCExecute can be restricted to a specific genomic region and can limit the number of generated scBAMs. An option is provided to automatically index the scBAMs files using samtools prior to executing the user-supplied command.

SCExecute is freely available (https://github.com/HorvathLab/NGS) as a self-contained binary package for 64-bit Linux and MacOS (Darwin) systems, and as Python3 source. The self-contained binary packages are appropriate for most Linux and MacOS users. The pythonic version requires pysam, numpy and scipy along with other packages (detailed instructions at https://github.com/HorvathLab/NGS).

#### Example of use

We demonstrate SCExecute using two variant callers - GATK and Strelka2 - run from a shell script with additional application of utilities for bam handling and variant filtering (SAMtools, BCFtools) (Van der Auwera *et al*., 2013; Kim *et al*., 2018; Li *et al*., 2009). We used publicly available scRNA-seq datasets generated on the 10xGenomics Chromium-3’UTR system on ten scRNA-seq datasets, including adrenal neuroblastoma, prostate cancer and MCF7 cell line, from 3 different studies (Dong *et al*., 2020; Ma *et al*., 2020; Ben-David *et al*., 2018). The data processing is shown on Figure 1a (See also the Supplementary Methods and Figures).

**Figure 1.**
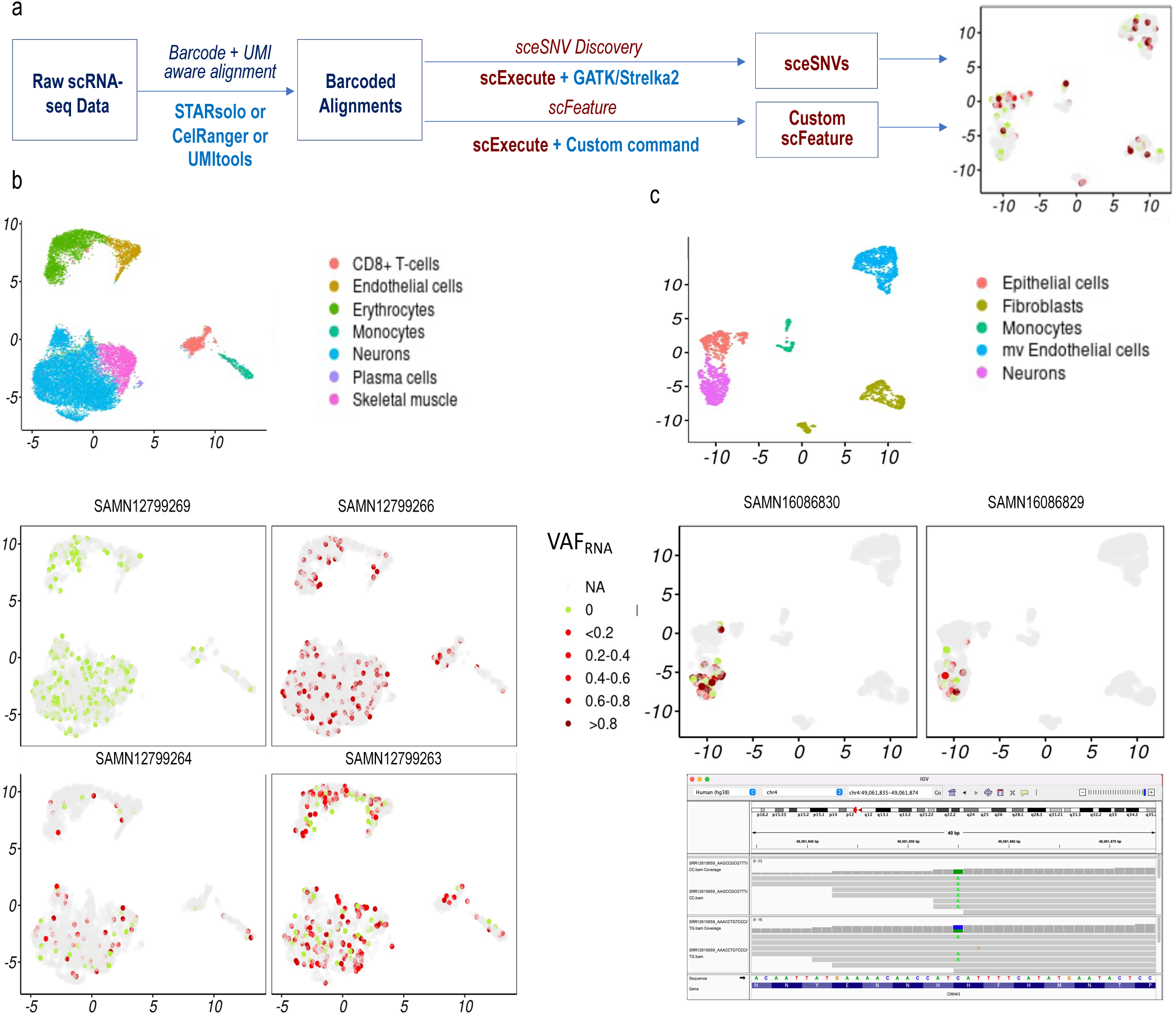
a. ScExecute data processing examples. b. UMAP projections showing cells classified by type (top) and visualizing the cell distribution and the cellular expressed variant allele frequency (VAF_RNA_) of the missense substitution rs4603 (1:151401549_T>C) in the gene *PSMB4* across the samples from the neuroblastoma dataset. The red color intensity shows the relative expression of the sceSNV in cells with at least 5 sequencing reads covering the SNV locus, and the green color indicates that all the reads covering the SNV locus carried the reference nucleotide (See Supplementary Methods). Cells in which the SNV locus is covered by less than 5 reads are shown in grey. The rs4603 VAF_RNA_ cell distribution is consistent with germline homozygous variant in sample SAMN12799266, heterozygous variant in samples SAMN12799264 and SAMN12799263, and absence in sample SAMN12799269. c. UMAP showing cells classified by type and the cell distribution and VAF_RNA_ of the missense substitution rs 1051447 (also reported as a somatic mutation, COSV56936745) in the gene CWH43 in the two prostate cancer samples (top). The CWH43 and COSV56936745 are mostly expressed in neurons. The IGV visualization of SNV positive cells shows mono-and bi-allelic expression of the SNV (bottom).

Briefly, the pooled scRNA-seq sequencing reads were aligned using the STARsolo module of STAR v.2.7.7a in 2-pass mode, with transcript annotations assembly GRCh38.79. Using SCExecute, variant calling was performed on the aligned reads of each cell, applying the HaplotypeCaller module of GATK v.4.2.0.0 and Strelka2 v.2.9.10 in parallel, followed by BCFtools variant quality filtering. This analysis identified up to 70K single cell-specific expressed Single Nucleotide Variants (sceSNVs) in two or more cells per sample (Figure 1b and c, and S_Table1). From those, between 11% and 36% sceSNVs per dataset were not previously reported in the Single Nucleotide Polymorphism Database (DbSNP, Sherry *et al*., 1999, Table_S2). For sceSNVs of interest, the corresponding scBAMs, optionally restricted to the sceSNV regions, can be saved using the scExecute “file template” option, and further explored, for example, through the Integrative Genomics Viewer (IGV, (Robinson *et al*., 2011), Figure 1c.

### Performance

The running time of SCExecute-based barcode-stratified analyses is driven largely by the execution time of the user-supplied command applied to each scBAM file. The primary execute-time attributable to SCExecute itself is the construction of the first batch of scBAMs – subsequent passes through the pooled scRNA-Seq BAM file are carried out in parallel with the execution of the user-supplied command and only as needed. For large enough batch sizes, applying the user-supplied command to each scBAM of the batch will take longer than each SCExecute pass through the pooled scRNA-Seq data. The SCExecute running time to create a batch of scBAMs is essentially invariant to the size of the batch and involves primarily I/O to access each alignment, extract the cellular barcode, and write out (some of) the reads. SCExecute will optionally use multiple processors to execute the user-supplied command in parallel. Total running time of a SCExecute-based analysis will therefore depend on the number of cellular barcodes, the execution time of the user-specified command, the number of processors used, and the time to create the first batch of scBAM files. Unless the user-specific command is very quick, the SCExecute specific contribution to running time is negligible.

For the genome-wide GATK variant call script applied to cellular-barcode stratified scBAMs, scRNA-Seq dataset SAMN09210328 contains 1858 cell-barcodes after alignment using STARsolo and filtering. The GATK variant calls script takes, on average, 3 minutes, per scBAM and the total execution time was 726 minutes using 8 processors, for an SCExecute running-time overhead of less than 5%.

## Discussion

Cell-level transcriptome analyses are essential to understand the details of each cell’s expressed features. To aid these analyses, we have developed SCExecute, which provides an efficient solution for custom cell-specific analyses from scRNA-seq data using existing tools, including such for bulk RNA- and DNA-sequencing data.

We demonstrate the SCEexecute application with variant callers designed for bulk sequencing data to identify sceSNVs. This analysis identified over 10,000 high-quality non-DbSNP SNVs across 10 datasets and 51,411 cells. SceSNVs from 10xGenomics scRNA-seq data are vastly understudied, as bulk-designed variant callers estimate quality metrics, including allele frequency and/or genotype confidence, based on all (pooled) scRNA-seq reads in a sample. As a result, SNVs with low allele frequency and/or uncertain genotypes in the pooled data can be filtered out. At the same time, it is well acknowledged that post-zygotically occurring SNVs (such as somatic and mosaic mutations), being present in only a proportion of cells, can result in low allele frequency and uncertain genotypes. Similar considerations apply to RNA-stemming variants, such as those resulting from RNA-editing or transcriptional infidelity. Importantly, the same reasoning is expected to apply to any cell-specific feature, including splicing and allele-specific expression. By bringing the estimations to the level of the cell SCExecute helps to expose the unique and specific cellular features and discover low (cellular) frequency transcriptome features.

## Supporting information

SCExecute_Supplementary_Materials

## Authors Contribution

NE and AH developed the concept, NE developed and implemented the software, CD, PNM, HL and NG tested and optimized the software and performed the analyses. AH devised and supervised the study and wrote the manuscript. All authors have read and approved the final manuscript.

