## Supplementary material for "SCExecute: cell barcode-stratified analyses of scRNA-seq data": SCExecute_Supplementary_Materials: Figure_S1.pdf

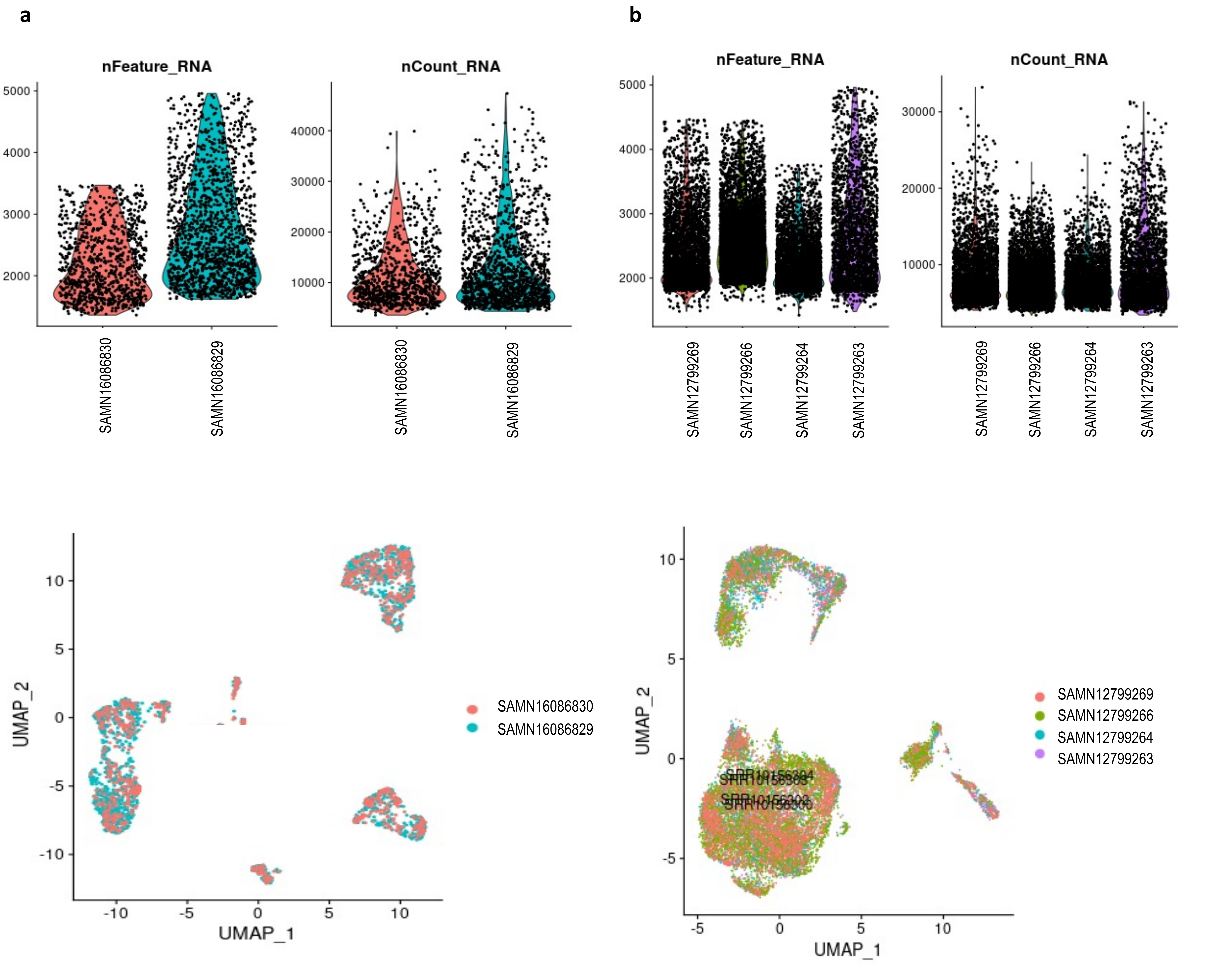


**Figure_S1**. Features (genes) and read conts distribution (top) after quality filtering and batch effect removal of the prostate cancer dataset (**a**) and the neuroblastoma dataset (**b**).
