## Supplementary material for "SCExecute: cell barcode-stratified analyses of scRNA-seq data": SCExecute_Supplementary_Materials: Figure_S2.pdf

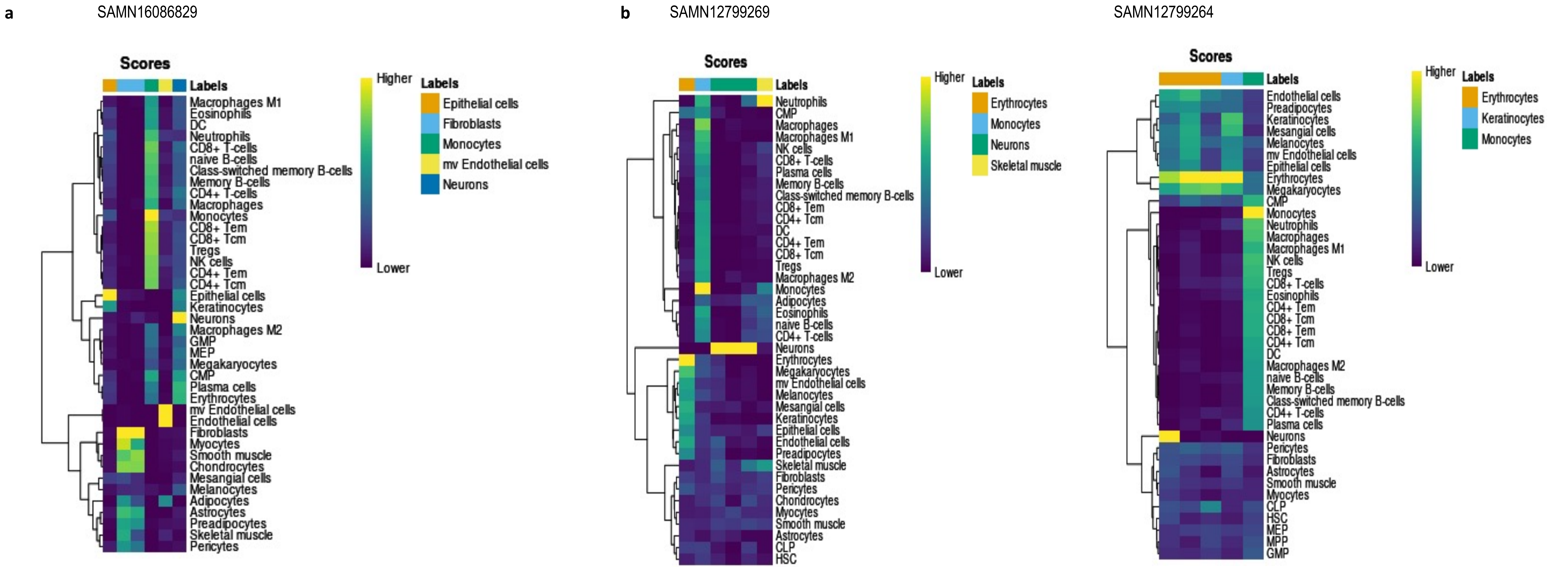


**Figure_S2**. Heatmaps showing cell-types similarity toknown cell types of the sample SAMN16086829 from the prostate cancer dataset (a) and samples SAMN12799269 and SAMN12799264 of the neuroblastoma dataset as determined by SingleR
