## Supplementary material for "SCExecute: cell barcode-stratified analyses of scRNA-seq data": SCExecute_Supplementary_Materials: Supplementary_Methods.pdf

*Sequencing Datasets*

To exemplify scExecute we utilized publicly available scRNA-seq data from three different cancer studies providing prostate cancer tissue (Ma *et al.*, 2020), neuroblastoma (Dong *et al.*, 2020) and the human breast carcinoma cell line MCF7 (Ben-David *et al.*, 2018). The experimental design and the data generation are described in detail in the original studies. All the studies utilized 10x Genomics Chromium Single Cell 3′ Workflow protocol for the libraries' generation, and the libraries were sequenced on an Illumina NextSeq 500 platform. The sequencing datasets were downloaded from the NCBI Sequence Read Archive (SRA) (accessed on 1 April 2021) under the accession numbers listed in Table_S1.

*Alignment and Variant call*

The pooled raw scRNA-seq sequencing reads were aligned using the STARsolo module of STAR v.2.7.7a in 2-pass mode, with transcript annotations assembly GRCh38.79 (Kaminow *et al.*, 2021). Transcriptome-wide variant calling was performed on each SCbam using SCExecute in conjunction with the HaplotypeCaller module of GATK v.4.2.0.0 and Strelka2 v.2.9.10 in parallel; both tools were used in their default setting (Van der Auwera *et al.*, 2013; Kim *et al.*, 2018). The GATK HaplotypeCaller was preceded by the assignment of read groups using the GATK module AddOrReplaceReadGroups, followed by splitting reads that contain Ns in their cigar string with the GATK module SplitNCigarReads.

For our analysis, we focused on the SNVs (i.e. indels and other variants were filtered out). The sceSNV calls from the individual alignments were filtered using the bcftools utility (v.1.10.2) of SAMtools retaining sceSNVs with QUAL (Phred-scaled probability) $>$ 100, MQ (mapping quality) $>$ 60, and QD (quality by depth) $>$ 2. SNV loci were annotated using SeattleSeq v.16.00 (dbSNP build 154), and SNVs positioned in non-repetitive regions were retained for further analysis. For sceSNVs of interest, the corresponding SCbams, optionally restricted to the sceSNV regions, can be saved using the scExecute "file template" option, and further explored, for example, through the Integrative Genomics Viewer (IGV, (Robinson *et al.*, 2011), Figure 1c.

*CQuality Assessments*

To define likely cell types, we used read-count matrices with the raw gene counts per cell generated by STARsolo. We normalized and scaled the expression data using the SCTransform function, as implemented in Seurat v.3.0 (Butler *et al.*, 2018). The cell-feature distributions were then plotted to identify and filter out the outliers and low-quality cells, which we defined after examination of the cell feature distribution (Supplementary Figure_S1). Specifically, based on the cell and feature distribution, we have filtered out: (1) cells with mitochondrial gene expression of above between 10% and 20%, (2) cells with fewer than 1000 genes, and (3) cells with more than between 3500 and 5500 detected genes (to remove potential doublets). The Seurat-processed gene expression values were also used to remove batch effects and cell cycle effects, as well as for cell type assessments.

*Cell Types Classifications*

To define likely cell types with known cell types, we used SingleR v.1.0.5 (Aran *et al.*, 2019), as previously described (Liu *et al.*, 2021; N. Prashant *et al.*, 2021; N. M. Prashant *et al.*, 2021). SingleR defines likely cell types, comparing genome-wide expression profile of each cell to a database of reference cells' whole transcriptome expression (BluePrint + ENCODE datasets). To select the expression profile corresponding to known cell type, the analysis is rerun iteratively with the top cell types from the previous step.

*$VAF_{RNA}$ Estimation and SNV Distribution Plotting*

Single-cell level VAFRNA was assessed from the pooled scRNA-seq alignments using scReadCounts v.1.1.4, as we have previously described (N. M. Prashant et al., 2021). When provided with barcoded scRNA-seq alignments and genomic loci of interest (with alleles), SCReadCounts tabulates the reference and variant read counts ($n_{ref}$ and $n_{var}$, respectively), and generates a cell-SNV matrix with the $VAF_{RNA}$ estimated at a user-defined threshold of the minimum number of required sequencing reads (minR) for a confident $VAF_{RNA}$ assessment. For the analysis presented herein, we used minR ≥ 5, which excludes from the estimation those positions covered by an insufficient number of reads (in this case 5). The cell-SNV $VAF_{RNA}$ matrix is than used as an input together with outputs of Seurat and SingleR to plot the SNV-distributions on the two-dimensional UMAP projections.
