## Supplementary material for "SCExecute: cell barcode-stratified analyses of scRNA-seq data": SCExecute_Supplementary_Materials: Table_S1_stats.pdf

Table S1. SceSNVs statistics and technical metrics in 10 scRNA-seq sets.

| Source | SampleID | seq read length | N cells | Mean Reads/ cell | Mean UMI/ cell | Total N sceSNVs in 2+ cells | non-DbSNP sceSNVs (novel) in 2+ cells | |
|---|---|---|---|---|---|---|---|---|
| | | | | | | | Total N | per cell |
| prostate cancer | SAMN16086830 | 150 | 1455 | 121811 | 14298 | 68797 | 22020 | 15,1 |
| | SAMN16086829 | 150 | 2019 | 80528 | 14810 | 59708 | 19451 | 9,6 |
| neuro-blastoma | SAMN12799269 | 150 | 6994 | 20892 | 6969 | 53872 | 19217 | 2,7 |
| | SAMN12799266 | 150 | 12448 | 13313 | 5742 | 45640 | 12653 | 1,0 |
| | SAMN12799264 | 150 | 16554 | 9140 | 4446 | 36877 | 9731 | 0,6 |
| | SAMN12799263 | 150 | 4273 | 28516 | 8512 | 44654 | 15263 | 3,6 |
| MCF7 | SAMN09210331 | 100 | 1749 | 37605 | 21682 | 7058 | 780 | 0,4 |
| | SAMN09210329 | 100 | 2778 | 20836 | 15222 | 3784 | 528 | 0,2 |
| | SAMN09210328 | 100 | 1891 | 38597 | 28034 | 3432 | 411 | 0,2 |
| | SAMN09210327 | 100 | 1250 | 52928 | 28223 | 5190 | 661 | 0,5 |
| **Statistics** | | | **sum 51411** | **mean 41249** | **mean 13302** | **sum 329012** | **sum 100715** | **mean 3,4** |
